# Appraising the causal relevance of DNA methylation for risk of lung cancer

**DOI:** 10.1101/287888

**Authors:** Thomas Battram, Rebecca C Richmond, Laura Baglietto, Philip C Haycock, Vittorio Perduca, Stig E Bojesen, Tom R Gaunt, Gibran Hemani, Florence Guida, Robert Carreras-Torres, Rayjean Hung, Christopher I Amos, Joshua R Freeman, Torkjel M Sandanger, Therese H Nøst, Børge Nordestgaard, Andrew E Teschendorff, Silvia Polidoro, Paolo Vineis, Gianluca Severi, Allison M Hodge, Graham G Giles, Kjell Grankvist, Mikael B Johansson, Mattias Johansson, George Davey Smith, Caroline L Relton

## Abstract

**Background:** DNA methylation changes in peripheral blood have recently been identified in relation to lung cancer risk. Some of these changes have been suggested to mediate part of the effect of smoking on lung cancer. However, limitations with conventional mediation analyses mean that the causal nature of these methylation changes has yet to be fully elucidated.

**Methods:** We first performed a meta-analysis of four epigenome-wide association studies (EWAS) of lung cancer (918 cases, 918 controls). Next, we conducted a two-sample Mendelian randomization analysis, using genetic instruments for methylation at CpG sites identified in the EWAS meta-analysis, and 29,863 cases and 55,586 controls from the TRICL-ILCCO lung cancer consortium, to appraise the possible causal role of methylation at these sites on lung cancer.

**Results:** 16 CpG sites were identified from the EWAS meta-analysis (FDR < 0.05), 14 of which we could identify genetic instruments for. Mendelian randomization provided little evidence that DNA methylation in peripheral blood at the 14 CpG sites play a causal role in lung cancer development (FDR>0.05), including for cg05575921-*AHRR* where methylation is strongly associated with both smoke exposure and lung cancer risk.

**Conclusions:** The results contrast with previous observational and mediation analysis, which have made strong claims regarding the causal role of DNA methylation. Thus, previous suggestions of a mediating role of methylation at sites identified in peripheral blood, such as cg05575921-*AHRR*, could be unfounded. However, this study does not preclude the possibility that differential DNA methylation at other sites is causally involved in lung cancer development, especially within lung tissue.

**Key Messages:**

- DNA methylation is a modifiable biomarker, giving it the potential to be targeted for intervention in many diseases, including lung cancer which is the most common cause of cancer-related death.
- This Mendelian randomization study attempted to evaluate whether there was a causal relationship, and thus potential for intervention, between DNA methylation measured in peripheral blood and lung cancer by assessing whether genetically altered DNA methylation levels impart differential lung cancer risks.
- Differential methylation at 14 CpG sites identified in epigenome-wide association analysis of lung cancer were assessed. Despite >99% power to detect the observational effect sizes, our Mendelian randomisation analysis gave little evidence that any of sites were causally linked to lung cancer.
- This is in stark contrast to previous analyses that suggested two CpG sites within the *AHRR* and *F2RL3* locus, which were also observed in this analysis, mediate >30% of the effect of smoking on lung cancer.
- Overall findings suggest there is little or no role of differential methylation at the CpG sites identified within the blood in the development of lung cancer. Thus, targeting these sites for prevention of lung cancer is unlikely to yield effective treatments.

## Background

Lung cancer is the most common cause of cancer-related death worldwide (1). Several DNA methylation changes have been recently identified in relation to lung cancer risk (2–4). Given the plasticity of epigenetic markers, any DNA methylation changes that are causally linked to lung cancer are potentially appealing targets for intervention (5, 6). However, these epigenetic markers are sensitive to reverse causation, being affected by cancer processes (6), and are also prone to confounding, for example by socio-economic and lifestyle factors (7, 8).

One CpG site, cg05575921 within the aryl hydrocarbon receptor repressor (*AHRR*) gene, has been consistently replicated in relation to both smoking (9) and lung cancer (2, 3, 10) and functional evidence suggests that this region could be causally involved in lung cancer (11). However, the observed association between methylation and lung cancer might simply reflect separate effects of smoking on lung cancer and DNA methylation, i.e. the association may be a result of confounding (12), including residual confounding after adjustment for self-reported smoking behaviour (13, 14). Furthermore, recent epigenome-wide association studies (EWAS) for lung cancer have revealed additional CpG sites which may be causally implicated in development of the disease (2, 3).

Mendelian randomization (MR) uses genetic variants associated with modifiable factors as instruments to infer causality between the modifiable factor and outcome, overcoming most unmeasured or residual confounding and reverse causation (15, 16). In order to infer causality, three core assumptions of MR should be met: 1) The instrument is associated with the exposure, 2) The instrument is not associated with any confounders, 3) The instrument is associated with the outcome only through the exposure. MR may be adapted to the setting of DNA methylation (17–19) with the use of single nucleotide polymorphisms (SNPs) that correlate with methylation of CpG sites, known as methylation quantitative trait loci (mQTLs) (20).

In this study, we performed a meta-analysis of four lung cancer EWAS (918 case-control pairs) from prospective cohort studies to identify CpG sites associated with lung cancer risk and applied MR to investigate whether the observed DNA methylation changes at these sites are causally linked to lung cancer.

## Methods

### EWAS Meta-analysis

We conducted a meta-analysis of four lung cancer case-control EWAS that assessed DNA methylation using the Illumina Infinium® HumanMethylation450 BeadChip. All EWAS are nested within prospective cohorts that measured DNA methylation in peripheral blood samples before diagnosis: EPIC-Italy (185 case-control pairs), Melbourne Collaborative Cohort Study (MCCS) (367 case-control pairs), Norwegian Women and Cancer (NOWAC) (132 case-control pairs) and the Northern Sweden Health and Disease Study (NSHDS) (234 case-control pairs). Study populations, laboratory methods, data pre-processing and quality control methods have been described in detail elsewhere (3) and outlined in the **Supplementary Methods.**

To quantify the association between the methylation level at each CpG and the risk of lung cancer we fitted conditional logistic regression models for beta values of methylation (which ranges from 0 (no cytosines methylated) to 1 (all cytosines methylated)) on lung cancer status for the four studies. The cases and controls in each study were matched, details of this are in the **Supplementary Methods.** Surrogate variables were computed in the four studies using the SVA R package (21) and the proportion of CD8+ and CD4+ T cells, B cells, monocytes, natural killer cells and granulocytes within whole blood were derived from DNA methylation (22). The following EWAS models were included in the meta-analysis: Model 1 – unadjusted; Model 2 – adjusted for 10 surrogate variables (SVs); Model 3 – adjusted for 10 SVs and derived cell proportions. EWAS stratified by smoking status was also conducted (never (N=304), former (N=648) and current smoking (N=857)). For Model 1 and Model 2, the case-control studies not matched on smoking status (EPIC-Italy and NOWAC) were adjusted for smoking.

We performed an inverse-variance weighted fixed effects meta-analysis of the EWAS (918 case-control pairs) using the METAL software (http://csg.sph.umich.edu/abecasis/metal/). Direction of effect, effect estimates and the I^2^ statistic were used to assess heterogeneity across the studies in addition to effect estimates across smoking strata (never, former and current). All sites identified at a false discovery rate (FDR)<0.05 in Model 2 and 3 were also present in the sites identified in Model 1. The effect size differences between models for all sites identified in Model 1 were assessed by a Kruskal-Wallis test and a post-hoc Dunn’s test. There was little evidence for a difference (P > 0.1), so to maximize inclusion into the MR analyses we took forward the sites identified in the unadjusted model (Model 1).

### Mendelian randomization

Two-sample MR was used to establish potential causal effect of differential methylation on lung cancer risk (23, 24). In the first sample, we identified mQTL-methylation effect estimates (β_GP_) for each CpG site of interest in an mQTL database from the Accessible Resource for Integrated Epigenomic Studies (ARIES) (http://www.mqtldb.org). Details on the methylation pre-processing, genotyping and quality control (QC) pipelines are outlined in the **Supplementary Methods.** In the second sample, we used summary data from a GWAS meta-analysis of lung cancer risk conducted by the Transdisciplinary Research in Cancer of the Lung and The International Lung Cancer Consortium (TRICL-ILCCO) (29,863 cases, 55,586 controls) to obtain mQTL-lung cancer estimates (β_GD_) (25).

For each independent mQTL (r^2^<0.01, we calculated the log odds ratio (OR) per SD unit increase in methylation by the formula β_GD_/β_GP_ (Wald ratio). Standard errors were approximated by the delta method (26). Where multiple independent mQTLs were available for one CpG site, these were combined in a fixed effects meta-analysis after weighting each ratio estimate by the inverse variance of their associations with the outcome. Heterogeneity in Wald ratios across mQTLs was estimated using Cochran’s Q test, which can be used to indicate horizontal pleiotropy (27). Differences between the observational and MR estimates were assessed using a Z-test for difference.

If there was evidence for an mQTL-CpG site association in ARIES in at least one time-point, we assessed whether the mQTL replicated across time points in ARIES (FDR < 0.05, same direction of effect). Further, we re-analysed this association using linear regression of methylation on each genotyped SNP available in an independent cohort (NSHDS), using rvtests (28) (**Supplementary Methods**). Replicated mQTLs were included where possible to reduce the effect of winner’s curse using effect estimates from ARIES. We assessed the instrument strength of the mQTLs by investigating the variance explained in methylation by each mQTL (r^2^) as well as the F-statistic in ARIES (**Supplementary Table 1**). The power to detect the observational effect estimates in the two-sample MR analysis was assessed *a priori*, based on an alpha of 0.05, sample size of 29,863 cases and 55,586 controls (from TRICL-ILCCO) and calculated variance explained (r^2^).

MR analyses were also performed to investigate the impact of methylation on lung cancer subtypes in TRICL-ILCCO: adenocarcinoma (11,245 cases, 54,619 controls), small cell carcinoma (2791 cases, 20,580 controls), and squamous cell carcinoma (7704 cases, 54,763 controls). We also assessed the association in never smokers (2303 cases, 6995 controls) and ever smokers (23,848 cases, 16,605 controls) (25). Differences between the smoking subgroups were assessed using a Z-test for difference.

We next investigated the extent to which the mQTLs at cancer-related CpGs were associated with four smoking behaviour traits which could confound the methylation-lung cancer association: number of cigarettes per day, smoking cessation rate, smoking initiation and age of smoking initiation using GWAS data from the Tobacco and Genetics (TAG) consortium (N=74,053) (29).

## Supplementary analyses

### Assessing the potential causal effect of AHRR methylation: one sample MR

Given previous findings implicating methylation at *AHRR* in relation to lung cancer (2, 3), we performed a one-sample MR analysis (30) of *AHRR* methylation on lung cancer incidence using individual-level data from the Copenhagen City Heart Study (CCHS) (357 incident cases, 8401 remaining free of lung cancer). Details of the phenotypic, methylation and genetic data, as well as the linked lung cancer data, are outlined in the **Supplementary Methods**.

An allele score of mQTLs located with 1Mb of cg05575921-*AHRR* was created and its association with *AHRR* methylation tested (**Supplementary Methods**). We investigated associations between the allele score and several potential confounding factors (sex, alcohol consumption, smoking status, occupational exposure to dust and/or welding fumes, passive smoking). We next performed MR analyses using two-stage Cox regression, with adjustment for age and sex, and further stratified by smoking status.

### Tumour and adjacent normal methylation patterns

DNA methylation data from lung cancer tissue and matched normal adjacent tissue (N=40 squamous cell carcinoma and N=29 adenocarcinoma), profiled as part of The Cancer Genome Atlas (TCGA), were used to assess tissue-specific DNA methylation changes across sites identified in the meta-analysis of EWAS, as outlined previously (31),

### mQTL association with gene expression

For the genes annotated to CpG sites identified in the lung cancer EWAS, we examined gene expression in whole blood and lung tissue using data from the gene-tissue expression (GTEx) consortium (32).

Analyses were conducted in Stata (version 14) and R (version 3.2.2). For the two-sample MR analysis we used the MR-Base R package TwoSampleMR (33). An adjusted P value that limited the FDR was calculated using the Benjamini-Hochberg method (34). All statistical tests were two-sided.

## Results

A flowchart representing our study design along with a summary of our results at each step is displayed in Figure 1.

**Figure 1.**
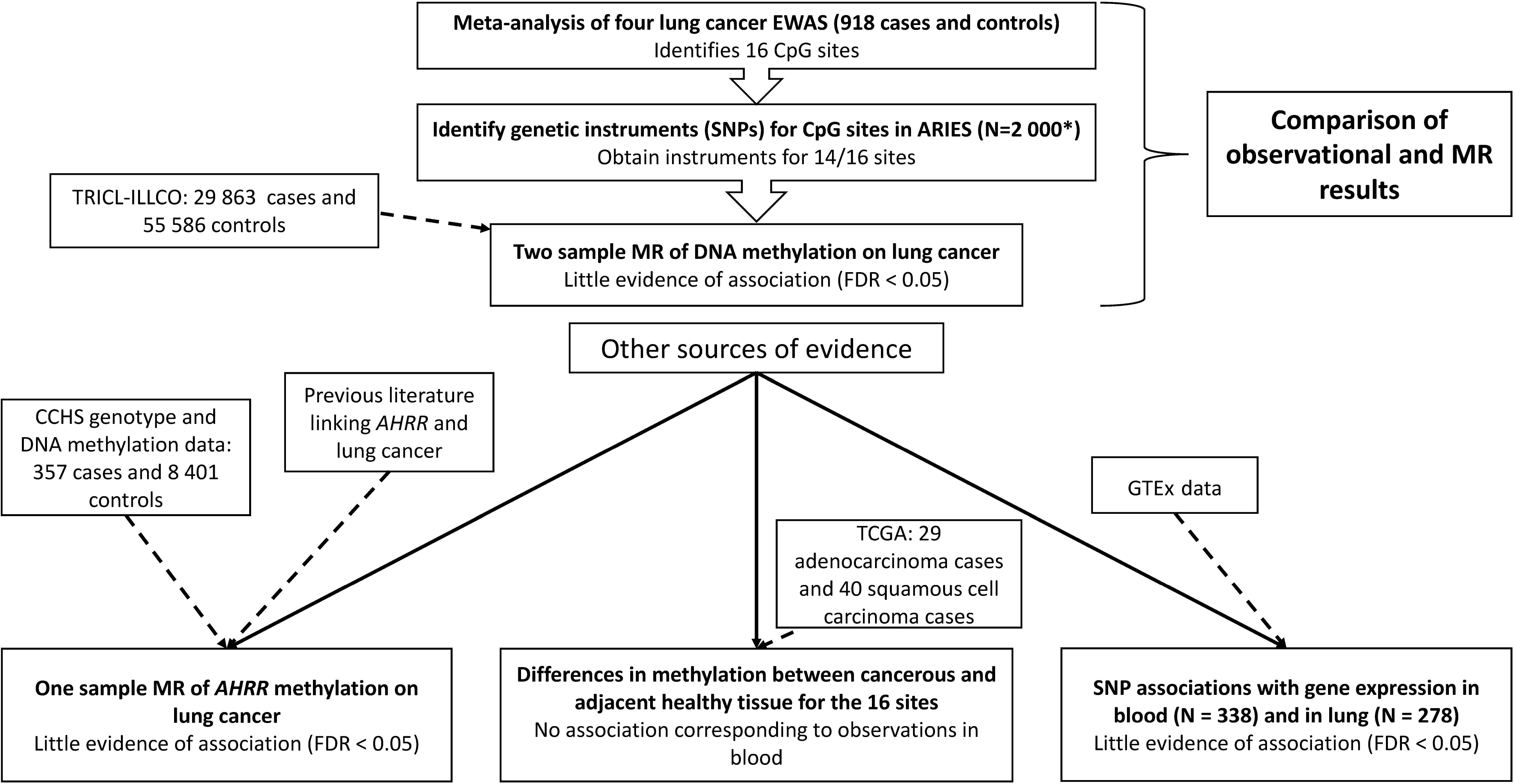
Study design with results summary. ARIES = Accessible Resource for Integrated Epigenomic Studies, TRICL-ILLCO = Transdisciplinary Research in Cancer of the Lung and The International Lung Cancer Consortium, MR = Mendelian randomization, CCHS = Copenhagen City Heart Study, TCGA = The Cancer Genome Atlas. * = 2 000 individuals with samples at multiple timepoints.

### EWAS meta-analysis

The basic meta-analysis adjusted for study-specific covariates identified 16 CpG sites which were hypomethylated in relation to lung cancer (FDR<0.05, Model 1, Figure 2). Adjusting for 10 surrogate variables (Model 2) and derived cell counts (Model 3) gave similar results (Table 1). The direction of effect at the 16 sites did not vary between studies (median I^2^=38.6) (**Supplementary Table 2**), but there was evidence for heterogeneity of effect estimates at some sites when stratifying individuals by smoking status (Table 1).

**Figure 2.**
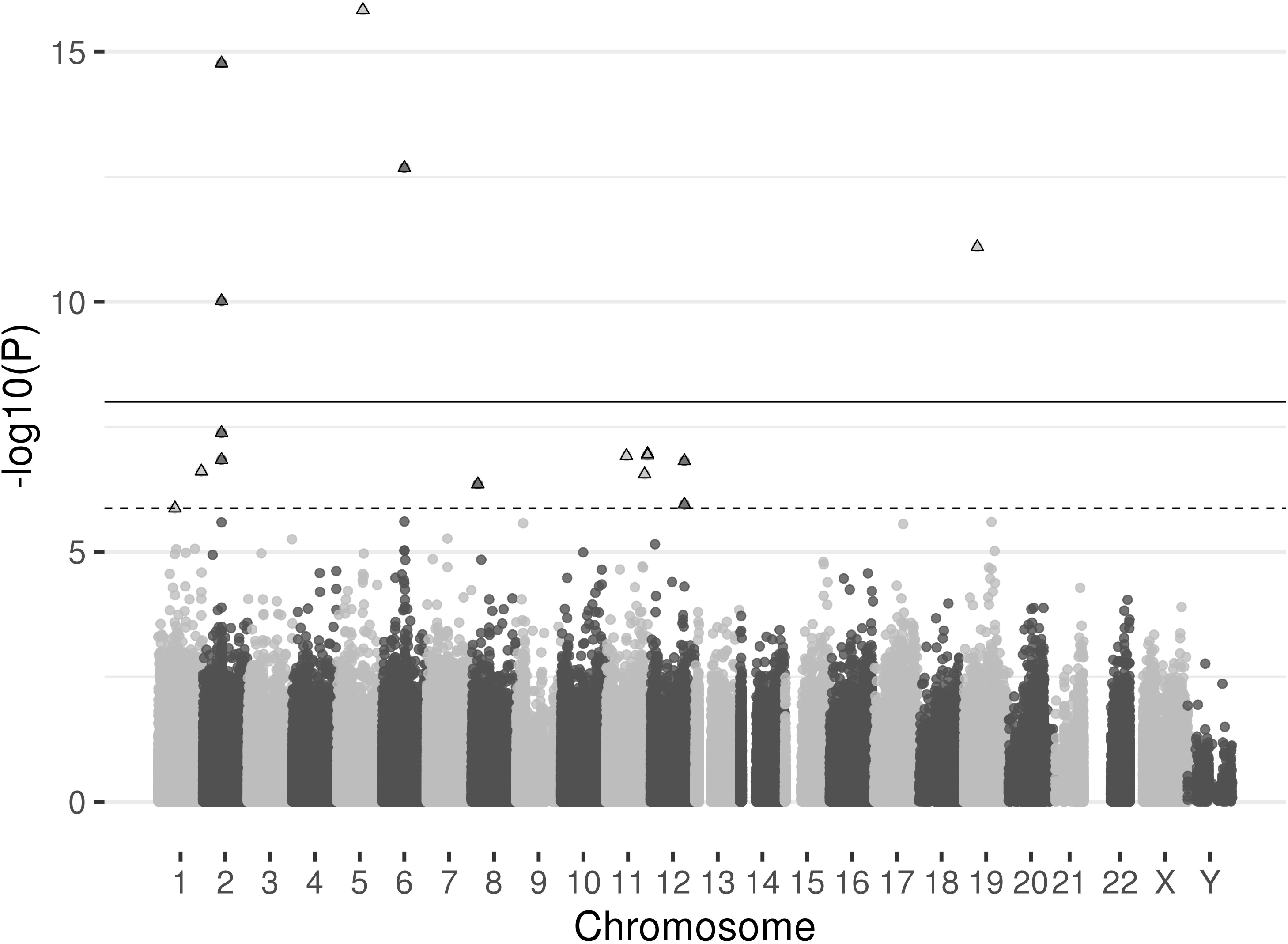
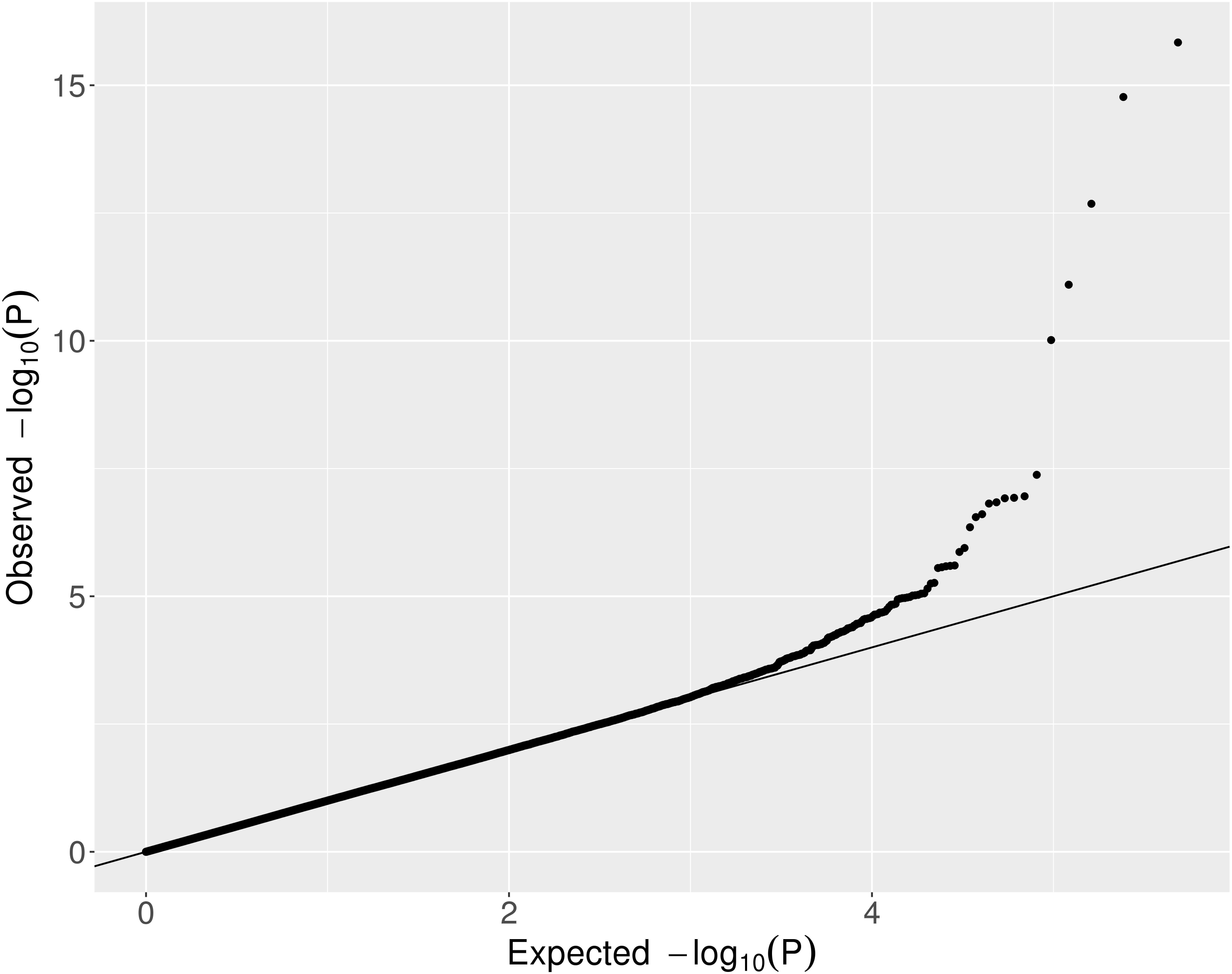
Observational associations of DNA methylation and lung cancer: A fixed effects meta-analysis of lung cancer EWAS weighted on the inverse variance was performed to establish the observational association between differential DNA methylation and lung cancer. **a)** Manhattan plot, all points above the solid line are at P < 1×10 and all points above the dashed line (and triangular points) are at FDR < 0.05. In total 16 CpG sites are associated with lung cancer (FDR < 0.05). **b)** Quantile-quantile plot of the EWAS results (same data as the Figure 2a Manhattan plot).

**Table 1.**
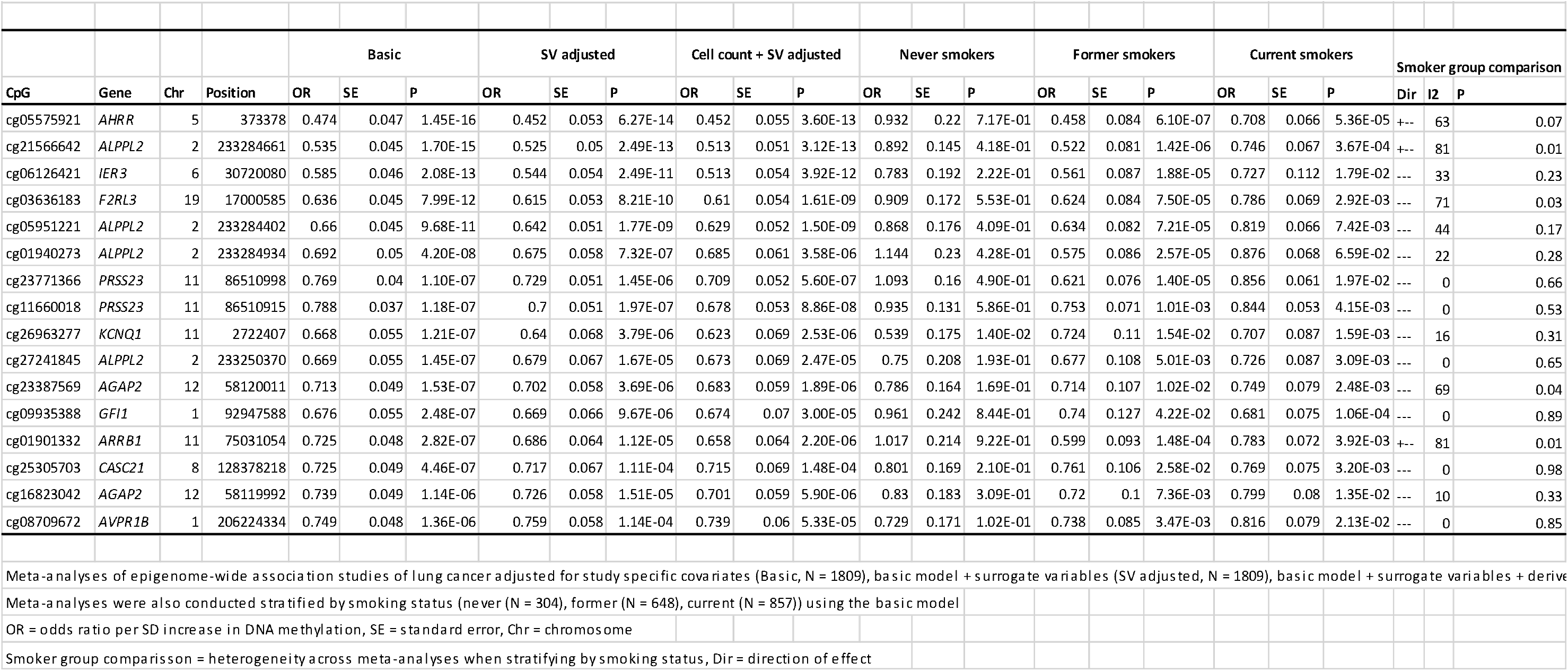
Meta-analyses of EWAS of lung cancer using four separate cohorts: 16 CpG sites associated with lung cancer at false discovery rate < 0.05.

### Mendelian randomization

We identified 15 independent mQTLs (r^2^<0.01) associated with methylation at 14 of 16 CpGs. Nine mQTLs replicated at FDR<0.05 in NSHDS (**Supplementary Table 3**). MR power analyses indicated >99% power to detect ORs for lung cancer of the same magnitude as those in the meta-analysis of EWAS.

There was little evidence for an effect of methylation at these 14 sites on lung cancer (FDR>0.05, **Supplementary Table 4**). For nine of 14 CpG sites the point estimates from the MR analysis were in the same direction as in the EWAS, but of a much smaller magnitude (Z-test for difference, P<0.001) (Figure 3).

**Figure 3.**
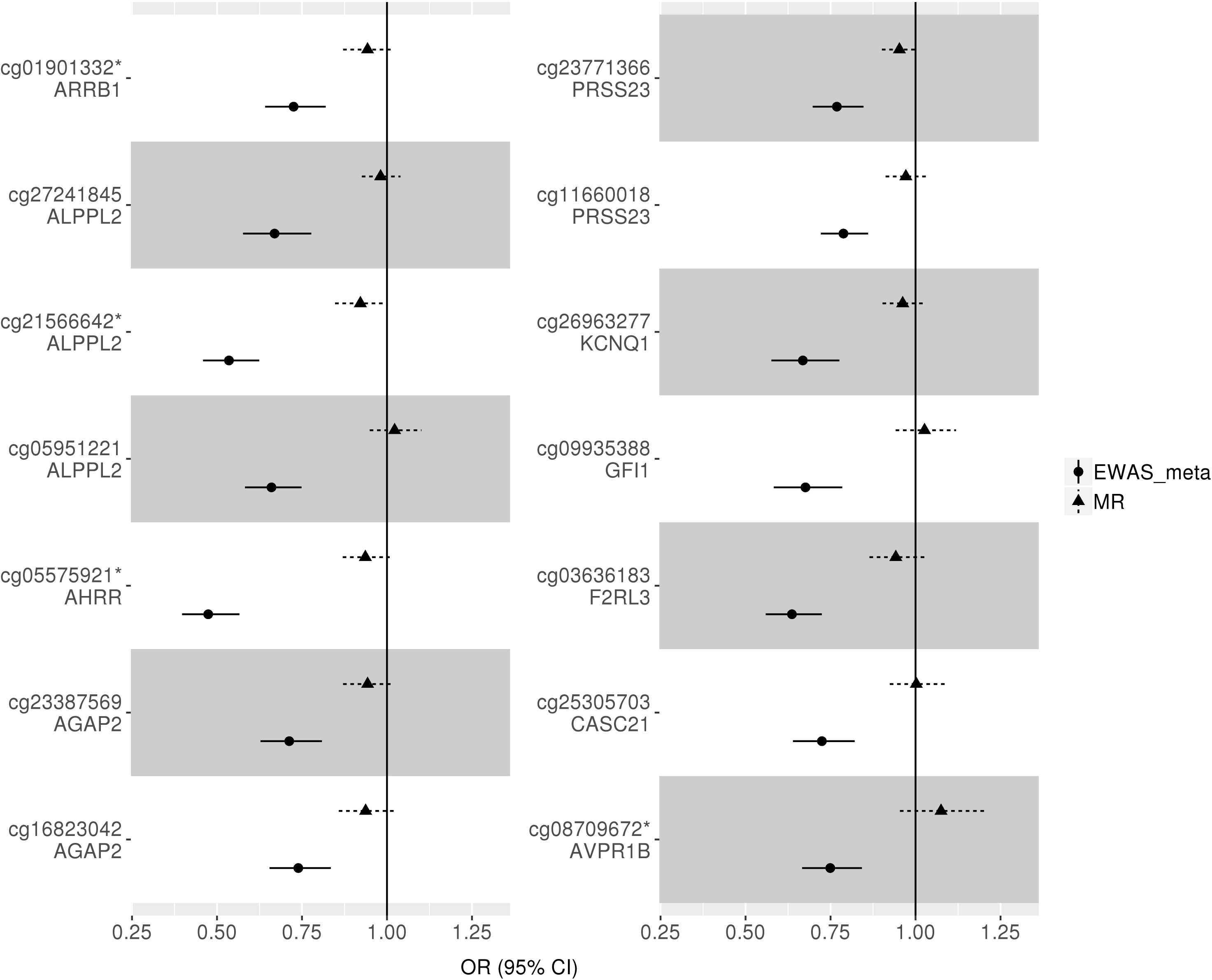
Mendelian randomization (MR) vs. observational analysis. Two-sample MR was carried out with methylation at 14/16 CpG sites identified in the EWAS meta-analysis as the exposure and lung cancer as the outcome. cg01901332 and cg05575921 had 2 instruments so the estimate was calculated using the inverse variance weighted method, for the rest the MR estimate was calculated using a Wald ratio. Only 14 of 16 sites could be instrumented using mQTLs from mqtldb.org. * = instrumental variable not replicated in independent dataset (NSHDS). The sites for which instrumental variables have not been replicated are cg01901332, cg21566642, cg05575921 and cg08709672. OR = odds ratio per SD increase in DNA methylation.

For 9 of out the 16 mQTL-CpG associations, there was strong replication across time points (**Supplementary Table 5**) and 9 out of 16 mQTL-CpG associations replicated at FDR<0.05 in an independent adult cohort (NSHDS). Using mQTL effect estimates from NSHDS for the nine CpG sites that replicated (FDR<0.05), findings were consistent with limited evidence for a causal effect of peripheral blood-derived DNA methylation on lung cancer (**Supplementary Figure 1**).

There was little evidence of different effect estimates between ever and never smokers at individual CpG sites (**Supplementary Figure 2**, Z-test for difference, P>0.5). There was some evidence for a possible effect of methylation at cg21566642-*ALPPL2* and cg23771366-*PRSS23* on squamous cell lung cancer (OR=0.85 [95% confidence interval (CI)=0.75,0.97] and 0.91 [95% CI=0.84,1.00] per SD [14.4% and 5.8%] increase, respectively) as well as methylation at cg23387569-*AGAP2*, cg16823042-*AGAP2*, and cg01901332-*ARRB1* on lung adenocarcinoma (OR=0.86 [95% CI=0.77,0.96], 0.84 [95% CI=0.74,0.95], and 0.89 [95% CI=0.80,1.00] per SD [9.47%, 8.35%, and 8.91%] increase, respectively). However, none of the results withstood multiple testing correction (FDR<0.05) (**Supplementary Figure 3**). For those CpGs where multiple mQTLs were used as instruments (cg05575921-*AHRR* and cg01901332-*ARRB1*), there was limited evidence for heterogeneity in MR effect estimates (Q-test, P>0.05, **Supplementary Table 6**).

Single mQTLs for cg05575921-*AHRR*, cg27241845-*ALPPL2*, and cg26963277-KCNQ1 showed some evidence of association with smoking cessation (former vs. current smokers), although these associations were not below the FDR<0.05 threshold (**Supplementary Figure 4**).

### Potential causal effect of *AHRR* methylation on lung cancer risk: one sample MR

In the CCHS, a per (average methylation-increasing) allele change in a four-mQTL allele score was associated with a 0.73% [95% CI=0.56,0.90] increase in methylation (P<1 × 10^−10^) and explained 0.8% of the variance in cg05575921-*AHRR* methylation (F-statistic=74.2). Confounding factors were not strongly associated with the genotypes in this cohort (P≥0.11) (**Supplementary Table 7**). Results provided some evidence for an effect of cg05575921 methylation on total lung cancer risk (HR=0.30 [95% CI=0.10,1.00] per SD (9.2%) increase) (**Supplementary Table 8**). The effect estimate did not change substantively when stratified by smoking status (**Supplementary Table 9**).

Given contrasting findings with the main MR analysis, where cg05575921-*AHRR* methylation was not causally implicated in lung cancer, and the lower power in the one-sample analysis to detect an effect of equivalent size to the observational results (power = 19% at alpha = 0.05), we performed further two-sample MR based on the four mQTLs using data from the TRICL-ILCCO consortium. Results showed no strong evidence for a causal effect of DNA methylation on total lung cancer risk (OR=1.00 [95% CI=0.83,1.10] per SD increase) (**Supplementary Figure 5**). There was also limited evidence for an effect of cg05575921-*AHRR* methylation when stratified by cancer subtype and smoking status (**Supplementary Figure 5**) and no strong evidence for heterogeneity of the mQTL effects (**Supplementary Table 9**). Conclusions were consistent when MR-Egger (27) was applied (**Supplementary Figure 5**) and when accounting for correlation structure between the mQTLs (**Supplementary Table 9**).

### Tumour and adjacent normal lung tissue methylation patterns

For cg05575921-*AHRR*, there was no strong evidence for differential methylation between adenocarcinoma tissue and adjacent healthy tissue (P=0.963), and weak evidence for hypermethylation in squamous cell carcinoma tissue (P=0.035) (Figure 4, **Supplementary Table 10**). For the other CpG sites there was evidence for a difference in DNA methylation between tumour and healthy adjacent tissue at several sites in both adenocarcinoma and squamous cell carcinoma, with consistent differences for CpG sites in *ALPPL2* (cg2156642, cg05951221 and cg01940273), as well as cg23771366-*PRSS23*, cg26963277-*KCNQ1*, cg09935388-*GFI1*, cg0101332-*ARRB1*, cg08709672-*AVPR1B* and cg25305703-*CASC21*. However, hypermethylation in tumour tissue was found for the majority of these sites, which is opposite to what was observed in the EWAS analysis.

**Figure 4.**
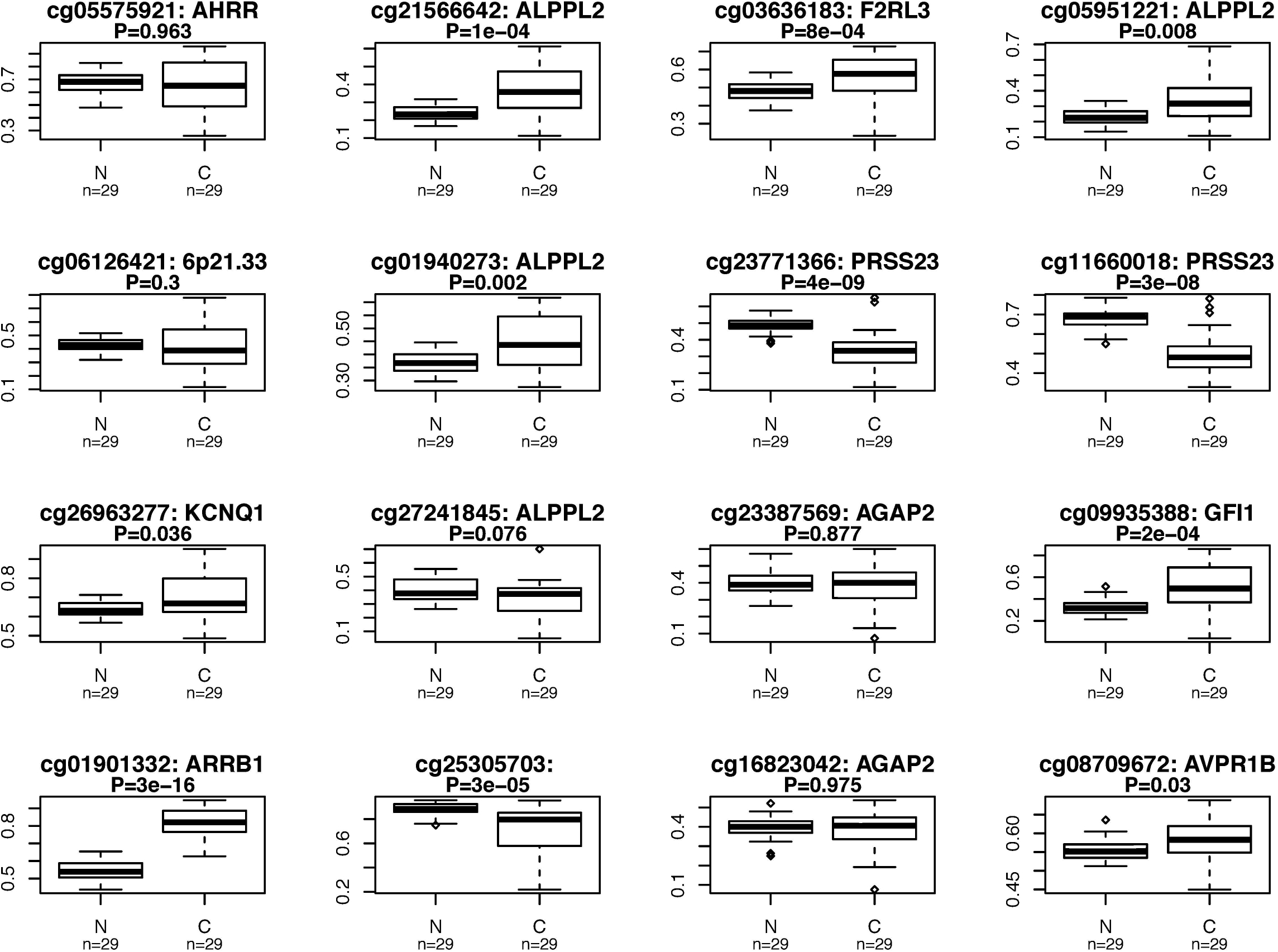
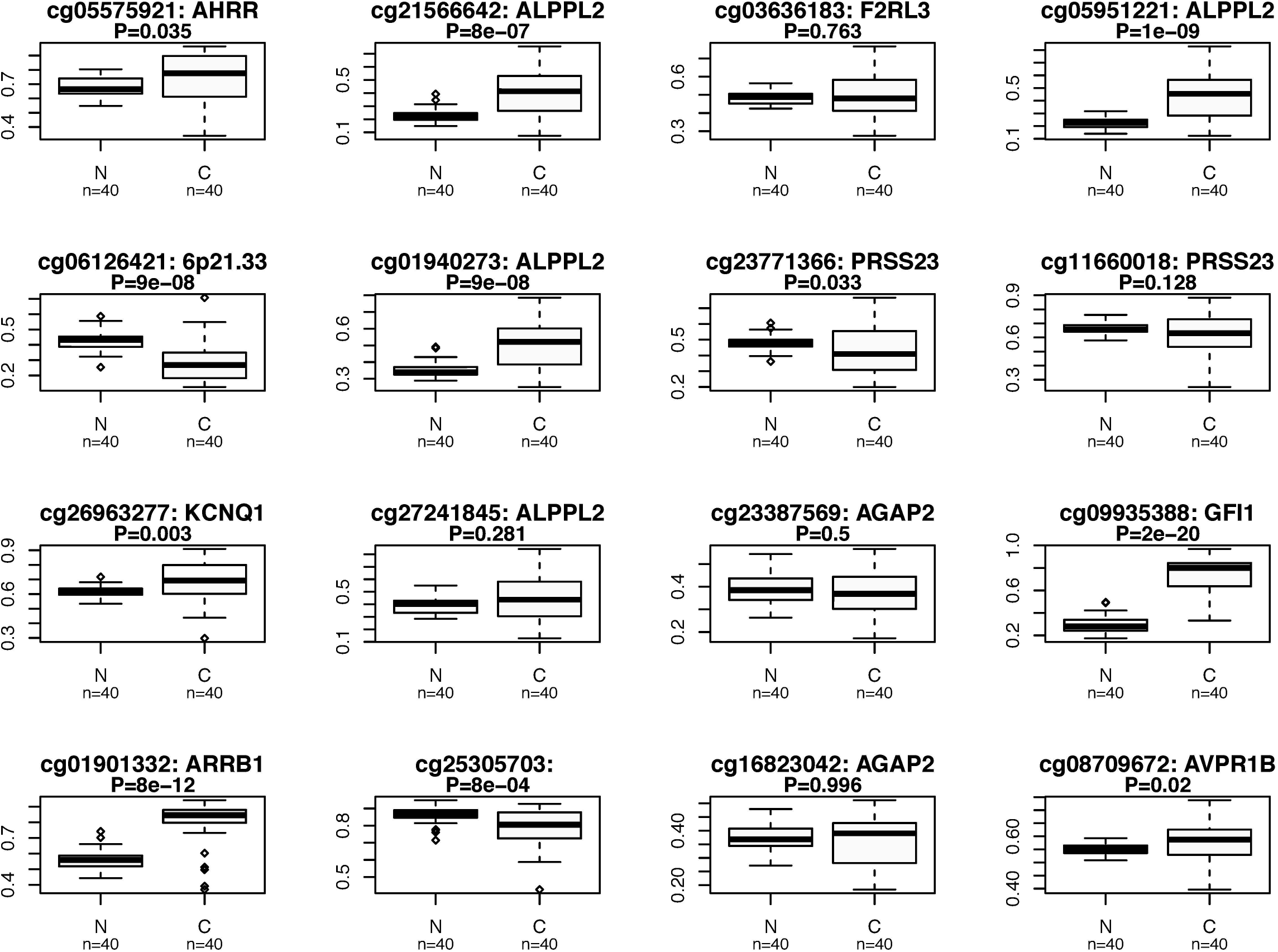
Differential DNA methylation in lung cancer tissue: A comparison of methylation at each of the 16 CpG sites identified in our meta-analysis was made between lung cancer tissue and adjacent healthy lung tissue for patients with a) lung adenocarcinoma and b) squamous cell lung cancer. Data from The Cancer Genome Atlas was used for this analysis.

### Gene expression associated with mQTLs in blood and lung tissue

Of the 10 genes annotated to the 14 CpG sites, eight genes were expressed sufficiently to be detected in lung (*AVPR1B* and *CASC21* were not) and seven in blood (*AVPR1B*, *CASC21* and *ALPPL2* were not). Of these, gene expression of *ARRB1* could not be investigated as the mQTLs in that region were not present in the GTEx data. rs3748971 and rs878481, mQTLs for cg21566642 and cg05951221 respectively, were associated with increased expression of *ALPPL2* (P=0.002 and P=0.0001). No other mQTLs were associated with expression of the annotated gene at a Bonferroni corrected P value threshold (P<0.05/19=0.0026) (**Supplementary Table 11**).

## Discussion

In this study, we identified 16 CpG sites associated with lung cancer, of which 14 have been previously identified in relation to smoke exposure (9) and six were highlighted in a previous study as being associated with lung cancer (3). This previous study used the same data from the four cohorts investigated here, but in a discovery and replication, rather than meta-analysis framework. Overall, using MR we found limited evidence supporting a potential causal effect of methylation at the CpG sites identified in peripheral blood on lung cancer. These findings are in contrast to previous analyses suggesting that methylation at two CpG sites investigated (in *AHRR* and *F2RL3*) mediated > 30% of the effect of smoking on lung cancer risk (2). This previous study used methods which are sensitive to residual confounding and measurement error that may have biased results (12, 35). These limitations are largely overcome using MR (12). While there was some evidence for an effect of methylation at some of the other CpG sites on risk of subtypes of lung cancer, these effects were not robust to multiple testing correction and were not validated in the analysis of tumour and adjacent normal lung tissue methylation nor in gene expression analysis.

A major strength of the study was the use of two-sample MR to integrate an extensive epigenetic resource and summary data from a large lung cancer GWAS to appraise causality of observational associations with >99% power. Evidence against the observational findings were also acquired through tissue-specific DNA methylation and gene expression analyses.

Limitations include potential “winner’s curse” which may bias causal estimates in a two-sample MR analysis towards the null if the discovery sample for identifying genetic instruments is used as the first sample, as was done for our main MR analysis using data from ARIES (36). However, findings were similar when using replicated mQTLs in NSHDS, indicating the potential impact of this bias was minimal (**Supplementary Figure 1**). Another limitation relates to the potential issue of consistency and validity of the instruments across the two samples. For a minority of the mQTL-CpG associations (4 out of 16), there was limited replication across time points and in particular, 6 mQTLs were not strongly associated with DNA methylation in adults. Caution is therefore warranted when interpreting the null results for the two-sample MR estimates for these CpG sites, which could be the result of weak-instrument bias.

The lack of independent mQTLs for each CpG site did not allow us to properly appraise horizontal pleiotropy in our MR analyses. Where possible we only included cis-acting mQTLs to minimise pleiotropy and investigated heterogeneity where there were multiple independent mQTLs. Three mQTLs were nominally associated with smoking phenotypes, but not to the extent that this would bias our MR results substantially. Some of the mQTLs used influence multiple CpGs in the same region, suggesting genomic control of methylation at a regional rather than single CpG level. This was untested, but methods to detect differentially methylated regions (DMRs) and identify genetic variants which proxy for them may be fruitful in probing the effect of methylation across gene regions.

A further limitation relates to the inconsistency in effect estimates between the one-and two-sample MR analysis to appraise the causal role of *AHRR* methylation. While findings in CCHS were supportive of a causal effect of *AHRR* methylation on lung cancer (HR=0.30 [95% CI=0.10,1.00] per SD), in two-sample MR this site was not causally implicated (OR=1.01 [95% CI=0.993, 1.03). We verified that this was not due to differences in the genetic instruments used, nor due to issues of weak instrument bias. Given the CCHS one-sample MR had little power (19% at alpha = 0.05) to detect a causal effect, we have more confidence in the results from the two-sample approach.

Peripheral blood may not be the ideal tissue to assess the association between DNA methylation and lung cancer. While a high degree of concordance in mQTLs has been observed across lung tissue, skin and peripheral blood DNA (37), we were unable to directly evaluate this here. A possible explanation for a lack of causal effect at *AHRR* is due to the limitation of tissue specificity as we found that the mQTLs used to instrument cg05575921 were not strongly related to expression of *AHRR* in lung tissue. However, findings from MR analysis were corroborated by the lack of evidence for differential methylation at *AHRR* between lung adenocarcinoma tissue and adjacent healthy tissue, and weak evidence for hypermethylation (opposite to the expected direction) in squamous cell lung cancer tissue. This result may be interesting in itself as smoking is hypothesized to influence squamous cell carcinoma more than adenocarcinoma. However, the result conflicts with that found in the MR analysis. Furthermore, another study investigating tumorous lung tissue (N=511) found only weak evidence for an association between smoking and cg05575921 *AHRR* methylation, that did not survive multiple testing correction (P=0.02) (38). However, our results do not fully exclude *AHRR* from involvement in the disease process. *AHRR* and AHR form a regulatory feedback loop, which means that the actual effect of differential methylation or differential expression of AHR/*AHRR* on pathway activity is complex (39). In addition, some of the CpG sites identified in the EWAS were found to be differentially methylated in the tumour and adjacent normal lung tissue comparison. While this could represent a false negative result of the MR analysis, it is of interest that differential methylation in the tissue comparison analysis was typically in the opposite direction to that observed in the EWAS. Furthermore, while this method can be used to minimize confounding, it does not fully eliminate the possibility of bias due to reverse causation (whereby cancer induces changes in DNA methylation) or intra-individual confounding e.g. by gene expression. Therefore, it doesn’t give conclusive evidence that DNA methylation changes at these sites are not relevant to the development of lung cancer.

While DNA methylation in peripheral blood may be predictive of lung cancer risk, according to the present analysis it is unlikely to play a causal role in lung carcinogenesis at the CpG sites investigated. Findings from this study issue caution over the use of traditional mediation analyses to implicate intermediate biomarkers (such as DNA methylation) in pathways linking an exposure with disease, given the potential for residual confounding in this context (12). However, the findings of this study do not preclude the possibility that other DNA methylation changes are causally related to lung cancer (or other smoking-associated disease) (40).

## Funding

This work was partly supported by a Wellcome Trust PhD studentship to TB [203746]; and by Cancer Research UK [C18281/A19169, C57854/A22171 and C52724/A20138]. This work was also supported by the UK Medical Research Council [MC_UU_12013/1 and MC_UU_12013/2], which funds a Unit at the University of Bristol where TB, RCR, PCH, TRG, GDS and CLR work. Funding to pay the Open Access publication charges for this article was provided by the University of Bristol RCUK.

The UK Medical Research Council and Wellcome [Grant ref: 102215/2/13/2] and the University of Bristol provide core support for ALSPAC. This publication is the work of the authors and TB, RCR and CLR will serve as guarantors for the contents of this paper.

Methylation data in the ALSPAC cohort were generated as part of the UK BBSRC funded [BB/I025751/1 and BB/I025263/1] Accessible Resource for Integrated Epigenomic Studies (ARIES, http://www.ariesepigenomics.org.uk).

## Competing interests statement

The authors declare no competing interests.

## Acknowledgements

For the contributions of ALSPAC data to our study: we are extremely grateful to all the families who took part, the midwives for their help in recruiting them, and the whole ALSPAC team, which includes interviewers, computer and laboratory technicians, clerical workers, research scientists, volunteers, managers, receptionists and nurses.

## References

1. Ferlay J, Soerjomataram I, Ervik M, Dikshit R, Eser S, C. M. GLOBOCAN 2012 v1.0, Cancer Incidence and Mortality Worldwide: IARC CancerBase No. 11 [Internet] 2013 [Available from: http://globocan.iarc.fr.

2. Fasanelli F, Baglietto L, Ponzi E, Guida F, Campanella G, Johansson M, et al. Hypomethylation of smoking-related genes is associated with future lung cancer in four prospective cohorts. Nat Commun. 2015;6:10192.

3. Baglietto L, Ponzi E, Haycock P, Hodge A, Bianca Assumma M, Jung CH, et al. DNA methylation changes measured in pre-diagnostic peripheral blood samples are associated with smoking and lung cancer risk. Int J Cancer. 2017;140(1):50–61.

4. McCarthy S, Das S, Kretzschmar W, Delaneau O, Wood AR, Teumer A, et al. A reference panel of 64,976 haplotypes for genotype imputation. Nat Genet. 2016;48(10):1279–83.

5. Strathdee G, Brown R. Aberrant DNA methylation in cancer: potential clinical interventions. Expert Rev Mol Med. 2002;4(4):1–17.

6. Jones PA, Baylin SB. The fundamental role of epigenetic events in cancer. Nat Rev Genet. 2002;3(6):415–28.

7. Borghol N, Suderman M, McArdle W, Racine A, Hallett M, Pembrey M, et al. Associations with early-life socio-economic position in adult DNA methylation. Int J Epidemiol. 2012;41(1):62–74.

8. Elliott HR, Tillin T, McArdle WL, Ho K, Duggirala A, Frayling TM, et al. Differences in smoking associated DNA methylation patterns in South Asians and Europeans. Clin Epigenetics. 2014;6(1):4.

9. Joehanes R, Just AC, Marioni RE, Pilling LC, Reynolds LM, Mandaviya PR, et al. Epigenetic Signatures of Cigarette Smoking. Circ Cardiovasc Genet. 2016;9(5):436–47.

10. Bojesen SE, Timpson N, Relton C, Davey Smith G, Nordestgaard BG. AHRR (cg05575921) hypomethylation marks smoking behaviour, morbidity and mortality. Thorax. 2017.

11. Zudaire E, Cuesta N, Murty V, Woodson K, Adarns L, Gonzalez N, et al. The aryl hydrocarbon receptor repressor is a putative tumor suppressor gene in multiple human cancers. J Clin Invest. 2008;118(2):640–50.

12. Richmond RC, Hemani G, Tilling K, Davey Smith G, Relton CL. Challenges and novel approaches for investigating molecular mediation. Hum Mol Genet. 2016;25(R2):R149–R56.

13. Fewell Z, Davey Smith G, Sterne JA. The impact of residual and unmeasured confounding in epidemiologic studies: a simulation study. Am J Epidemiol. 2007;166(6):646–55.

14. Munafo MR, Timofeeva MN, Morris RW, Prieto-Merino D, Sattar N, Brennan P, et al. Association Between Genetic Variants on Chromosome 15q25 Locus and Objective Measures of Tobacco Exposure. Jnci-J Natl Cancer I. 2012;104(10):740–8.

15. Davey Smith G, Hemani G. Mendelian randomization: genetic anchors for causal inference in epidemiological studies. Hum Mol Genet. 2014;23(R1):R89–98.

16. Davey Smith G, Ebrahim S. ’Mendelian randomization’: can genetic epidemiology contribute to understanding environmental determinants of disease? International Journal of Epidemiology. 2003;32(1):1–22.

17. Relton CL, Davey Smith G. Two-step epigenetic Mendelian randomization: a strategy for establishing the causal role of epigenetic processes in pathways to disease. Int J Epidemiol. 2012;41(1):161–76.

18. Relton CL, Davey Smith G. Mendelian randomization: applications and limitations in epigenetic studies. Epigenomics. 2015;7(8):1239–43.

19. Richardson TG, Zheng J, Davey Smith G, Timpson NJ, Gaunt TR, Relton CL, et al. Mendelian Randomization Analysis Identifies CpG Sites as Putative Mediators for Genetic Influences on Cardiovascular Disease Risk. Am J Hum Genet. 2017;101(4):590–602.

20. Gaunt TR, Shihab HA, Hemani G, Min JL, Woodward G, Lyttleton O, et al. Systematic identification of genetic influences on methylation across the human life course. Genome Biol. 2016;17:61.

21. Leek JT, Johnson WE, Parker HS, Fertig EJ, Jaffe AE, Storey JD. sva: Surrogate Variable Analysis. R package version 30. 2017.

22. Houseman EA, Accomando WP, Koestler DC, Christensen BC, Marsit CJ, Nelson HH, et al. DNA methylation arrays as surrogate measures of cell mixture distribution. Bmc Bioinformatics. 2012;13.

23. Inoue A, Solon G. Two-Sample Instrumental Variables Estimators. Rev Econ Stat. 2010;92(3):557–61.

24. Pierce BL, Burgess S. Efficient design for Mendelian randomization studies: subsample and 2-sample instrumental variable estimators. Am J Epidemiol. 2013;178(7):1177–84.

25. McKay JD, Hung RJ, Han Y, Zong X, Carreras-Torres R, Christiani DC, et al. Large-scale association analysis identifies new lung cancer susceptibility loci and heterogeneity in genetic susceptibility across histological subtypes. Nat Genet. 2017.

26. Thomas DC, Lawlor DA, Thompson JR. Re: Estimation of bias in nongenetic observational studies using “Mendelian triangulation” by bautista et al. Ann Epidemiol. 2007;17(7):511–3.

27. Bowden J, Davey Smith G, Burgess S. Mendelian randomization with invalid instruments: effect estimation and bias detection through Egger regression. International Journal of Epidemiology. 2015;44(2):512–25.

28. Zhan X, Hu Y, Li B, Abecasis GR, Liu DJ. RVTESTS: an efficient and comprehensive tool for rare variant association analysis using sequence data. Bioinformatics. 2016;32(9):1423–6.

29. Tobacco, Genetics C. Genome-wide meta-analyses identify multiple loci associated with smoking behavior. Nat Genet. 2010;42(5):441–7.

30. Haycock PC, Burgess S, Wade KH, Bowden J, Relton C, Davey Smith G. Best (but oft-forgotten) practices: the design, analysis, and interpretation of Mendelian randomization studies. Am J Clin Nutr. 2016;103(4):965–78.

31. Teschendorff AE, Yang Z, Wong A, Pipinikas CP, Jiao Y, Jones A, et al. Correlation of Smoking-Associated DNA Methylation Changes in Buccal Cells With DNA Methylation Changes in Epithelial Cancer. JAMA Oncol. 2015;1(4):476–85.

32. Consortium GT. The Genotype-Tissue Expression (GTEx) project. Nat Genet. 2013;45(6):580–5.

33. Hemani G, Zheng J, Wade KH, Elsworth B, Langdon R, Burgess S, et al. MR-Base: a platform for Mendelian randomization using summary data from genome-wide association studies. eLife. 2018;7:e34408.

34. Benjamini Y, Hochberg Y. Controlling the False Discovery Rate - a Practical and Powerful Approach to Multiple Testing. J Roy Stat Soc B Met. 1995;57(1):289–300.

35. Hemani G, Tilling K, Davey Smith G. Orienting the causal relationship between imprecisely measured traits using GWAS summary data. PLoS Genet. 2017;13(11):e1007081.

36. Burgess S, Thompson SG, Collaboration CCG. Avoiding bias from weak instruments in Mendelian randomization studies. International Journal of Epidemiology. 2011;40(3):755–64.

37. Shi J, Marconett CN, Duan J, Hyland PL, Li P, Wang Z, et al. Characterizing the genetic basis of methylome diversity in histologically normal human lung tissue. Nat Commun. 2014;5:3365.

38. Freeman JR, Chu S, Hsu T, Huang YT. Epigenome-wide association study of smoking and DNA methylation in non-small cell lung neoplasms. Oncotarget. 2016;7(43):69579–91.

39. Chen YT, Widschwendter M, Teschendorff AE. Systems-epigenomics inference of transcription factor activity implicates aryl-hydrocarbon-receptor inactivation as a key event in lung cancer development. Genome Biology. 2017;18.

40. Gao X, Zhang Y, Breitling LP, Brenner H. Tobacco smoking and methylation of genes related to lung cancer development. Oncotarget. 2016;7(37):59017–28.

